## Supplemental Files for "Multiomic ALS signatures highlight sex differences and molecular subclusters and identify the MAPK pathway as therapeutic target" (from another version of this preprint: version 2, 2024-02-07)

### List of supplemental figures


| No. | Title | page |
|---|---|---|
| 1 | Mapped entities | 2 |
| 2 | Sample distance human transcriptomics data | 3 |
| 3 | Kaplan-Meier curves for ALS mouse models and PCA analyses (transcriptomics) | 4 |
| 4 | Sample quality | 5 |
| 5 | Illustration of the KEGG pathway “Pathways of neurodegeneration - multiple diseases (hsa05022ID)” | 7 |
| 6 | Gene expression of genes involved in ALS subclusters by Tam et al. 2019 | 8 |
| 7 | Projection of transcriptomic subclusters onto proteomic data | 9 |
| 8 | Triplet-plots showing miRNA - mRNA - protein networks in human males and females | 10 |
| 9 | Density plot for factors identified with MOFA | 11 |
| 10 | Overview of MOFA factor 1 for male ALS patient samples | 11 |
| 11 | Protein-protein-interaction network for the interactors of MAPK1 among the target genes of miR-1273f | 12 |
| 12 | Overview of MOFA factor 12 for female ALS patient samples | 13 |
| 13 | Differential alternative splicing (DAS) analysis of the mouse models | 14 |
| 14 | Cell-type deconvolution | 15 |
| 15 | Summary of WGCNA for mouse models | 17 |
| 16 | Differential expression of proteins in mouse models | 19 |
| 17 | REVIGO clustering and sample correlation for mouse proteomics | 20 |
| 18 | Quadruple plots showing miRNA - mRNA - protein - phosphoprotein networks in the C9orf72 model | 21 |
| 19 | Differential expression of miRNA in mouse models | 22 |
| 20 | Differential regulation of MAPK pathways in human ALS patient subclusters | 23 |
| 21 | Western blot analysis of trametinib effects on MEK2, ERK1/2 and phospho-MEK2 with and without glutamate treatment in primary neuronal cultures | 24 |
| 22 | Western blot experiments - full membranes | 25 |
| 23 | Phospho-proteomics experiments for SOD1 mice treated with trametinib | 26 |
| 24 | Isolation of prefrontal cortex from ALS mouse models | 27 |

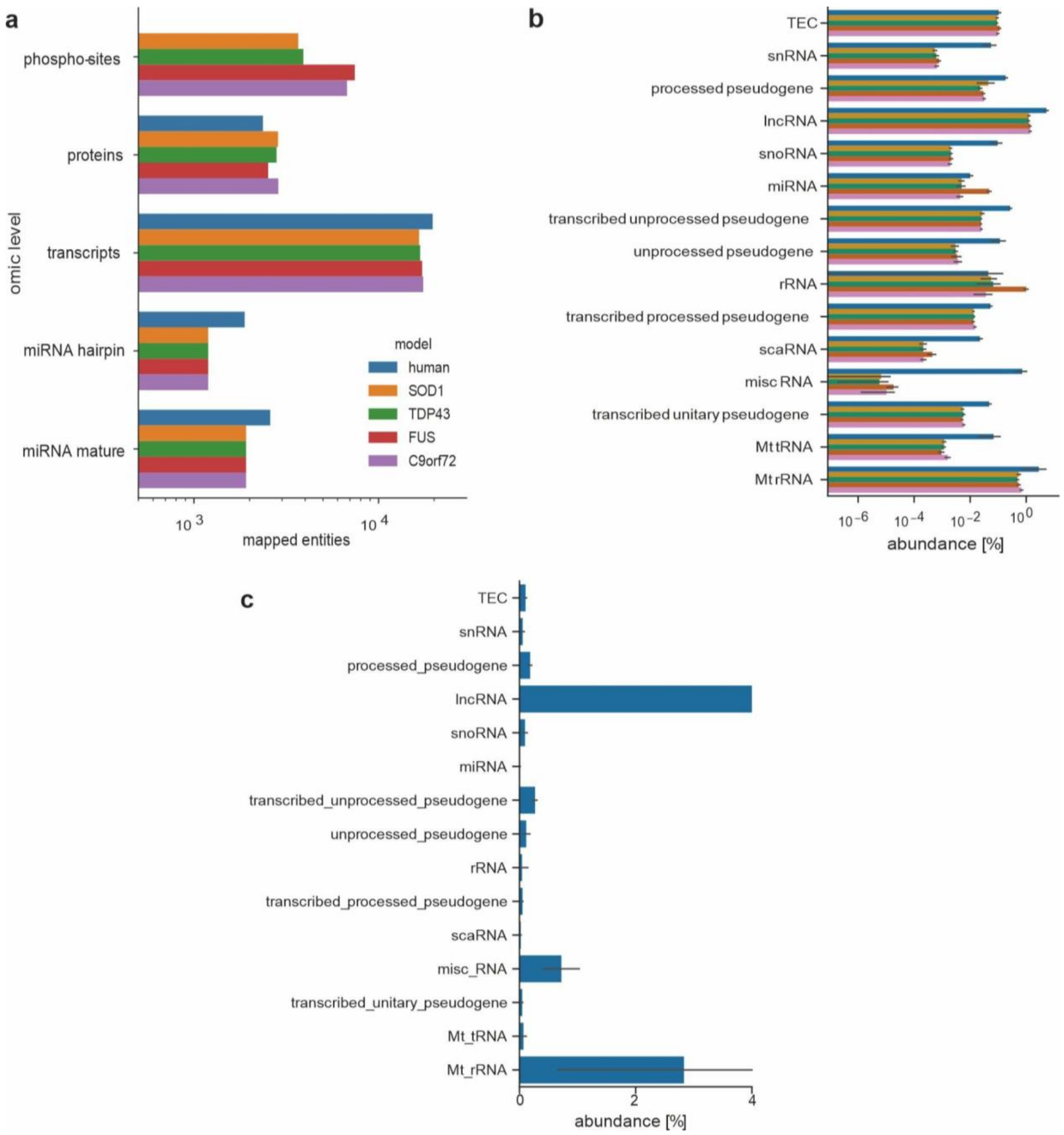


**Supplementary Fig. 1 Mapped entities.**

**a.** Barplot showing the number of mapped entities in phospho-sites, proteins, mRNA (transcripts), mature miRNA and hairpin miRNA for human samples and four mouse models (TDP43, FUS, C9orf72, and SOD1). **b.** Barplot showing the comparative abundance of mapped RNA types based on the mRNA sequencing for human and mouse samples, excluding protein-coding RNA. **c.** Barplot showing the abundance of mapped RNA types based on the mRNA sequencing only for humans, excluding protein-coding RNA.

TEC, to be experimentally confirmed; snRNA, small nuclear RNA; lncRNA, long non-coding RNA; snoRNA, small nucleolar RNA; miRNA, microRNA; rRNA, ribosomal RNA; scaRNA, small Cajal body-specific RNA; miscRNA, miscellaneous RNA; Mt tRNA, mitochondrial transfer RNA; Mt rRNA, mitochondrial ribosomal RNA.

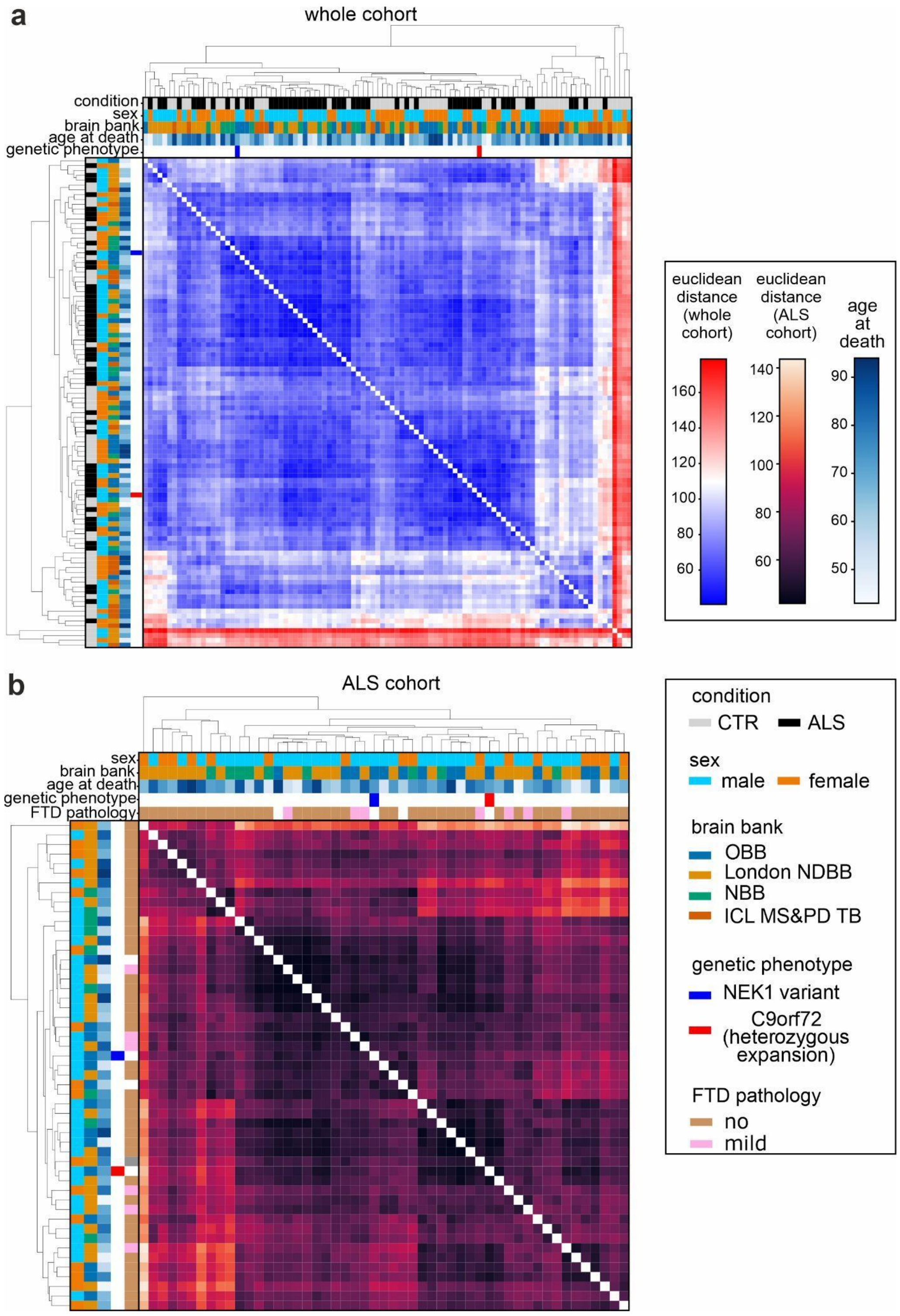


**Supplementary Fig. 2 Sample distance human transcriptomics data.**

Heatmaps showing the Euclidean distance between all samples based on the vst-transformed transcriptomics data for the whole human cohort (**a**) and the ALS cohort only (**b**). High values indicate a big difference between the samples. Condition, sex, brain bank, age at death and genetic phenotype of each sample are indicated. Samples are clustered according to their distance using hierarchical clustering. The diagonal (self-distance) is zero and masked from the color scale to allow a better distinction of the other sample distances. ICL MS&PD TB: Imperial College London - Multiple Sclerosis and Parkinson's Tissue Bank; NBB: The Netherlands Brain Bank; OBB: Oxford Brain Bank; London NDBB: London Neurodegenerative Diseases Brain Bank.

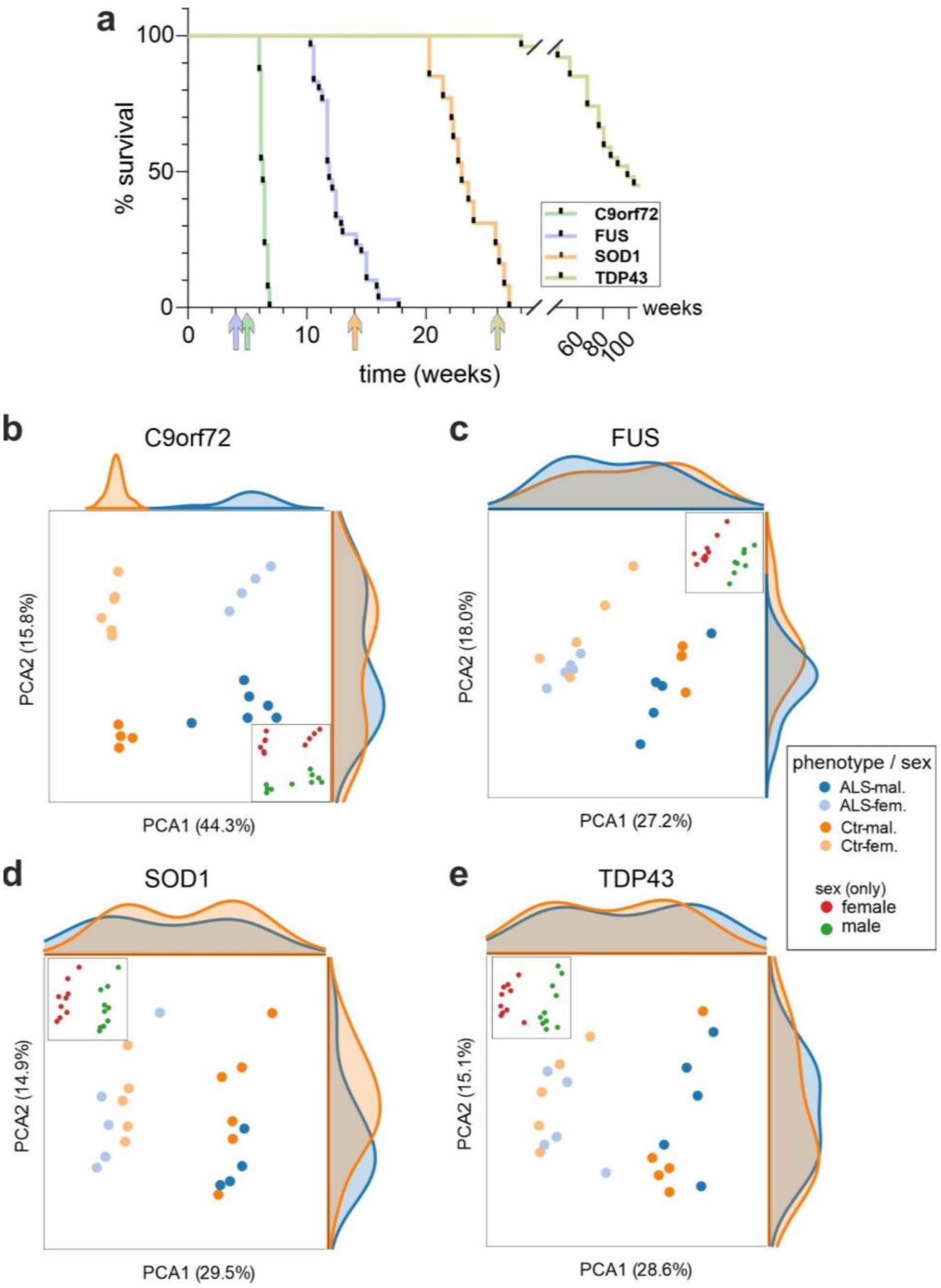


**Supplementary Fig. 3 Kaplan-Meier curves for ALS mouse models and PCA analyses (transcriptomics).**

**a.** Expected survival depicted in Kaplan-Meier curves for each of the selected ALS mouse models. Arrows indicate the time point of sacrifice for each mouse model (FUS=4 weeks; C9orf72=4.5 weeks; SOD1=14 weeks; TDP43=26 weeks). Animals were either pre-symptomatic (SOD1, FUS, TDP43) or showed early motor symptoms (C9orf72) at the time of sacrifice. **b-e.** Principal component analysis (PCA) on the 500 most variable genes of each of the mouse models: C9orf72 **(b)**; FUS **(c)**; SOD1 **(d)**; TDP43 **(e)**. Sex separation is indicated in red (females) and green (males).

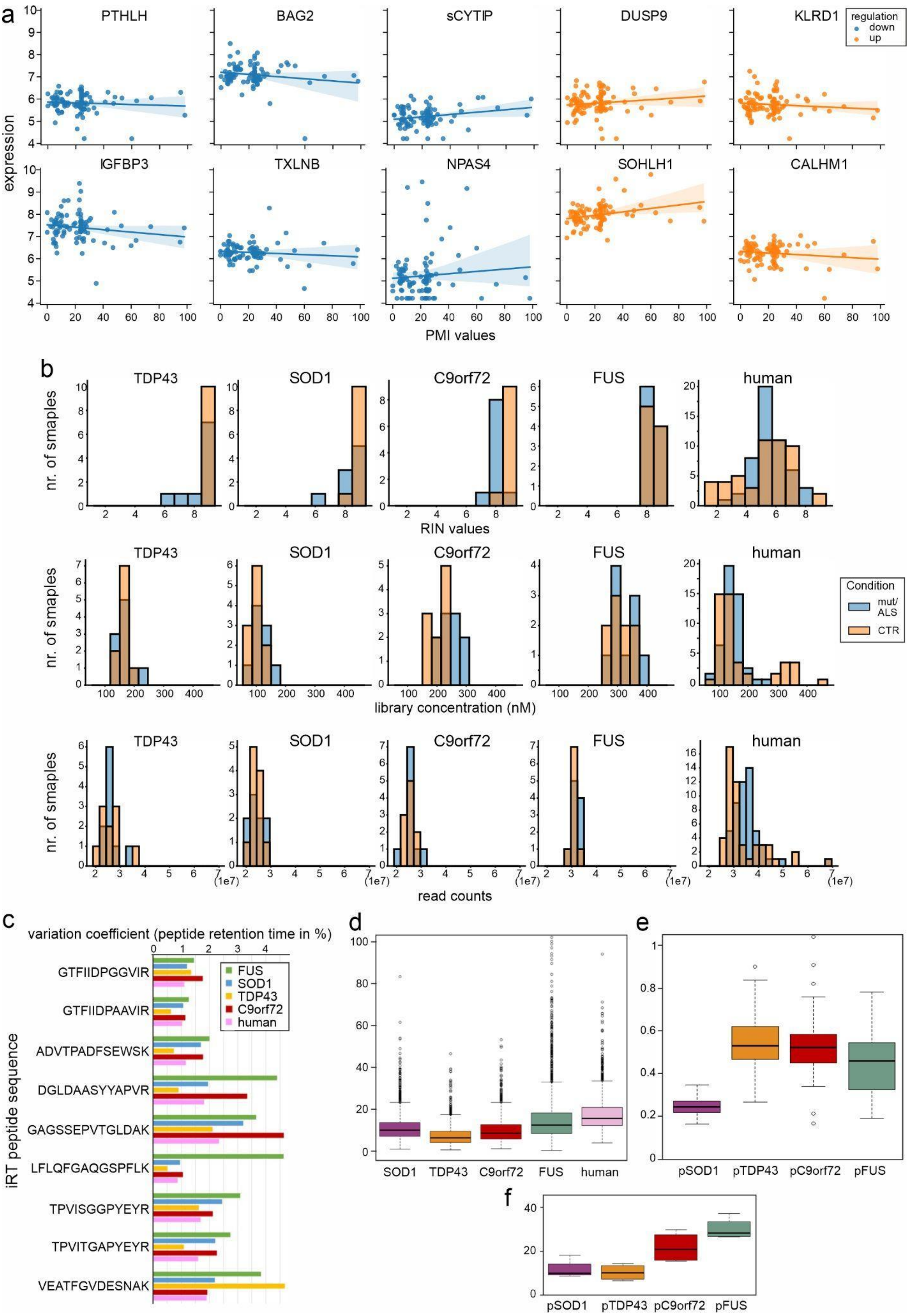


**Supplementary Fig. 4 Sample quality.**

**a.** Linear regression plots showing the correlation between postmortem interval (PMI) and the expression of selected neuronal marker genes in human brain samples (downregulated in blue and upregulated in orange. X-axis, PMI; y-axis, expression in normalized number of transcripts. PMI had no significant effect on the expression of these neuronal markers. **b.** Barplots showing the RNA integrity number (RIN), library concentration, read count and RNA counts for control (orange) and ALS (blue) samples. X-axis, RIN or library concentration or read counts; y-axis, the number of samples. **c.** External quality control for the proteomics analysis: coefficient of variation (in percentage) calculated on the retention time values of the nine iRT synthetic peptides that were observed in each proteomic cohort. **d.** Internal quality control for the proteomics analysis: boxplots representing the coefficients of variation of the LFQ intensities of all pool QC samples for each proteomic cohort. Those pool QC samples are a combination of equal amounts of each sample and underwent the same sample preparation steps as every other sample. Even for large cohorts of samples, the median CV <20%, highlighting the stability of the LC-MS/MS system. **e.** Quality controls for the phosphoproteomics cohorts: boxplots of the ratio (light phosphomix intensity/heavy phosphomix intensity) for each phosphoproteomic cohort. This allows us to evaluate the efficiency of the phosphopeptides enrichment step. **f.** Quality controls for the phosphoproteomics cohorts: boxplot of the CVs on the ratios (dividing (light phosphomix intensity) by (heavy phosphomix intensity)) for each phosphoproteomic cohort.

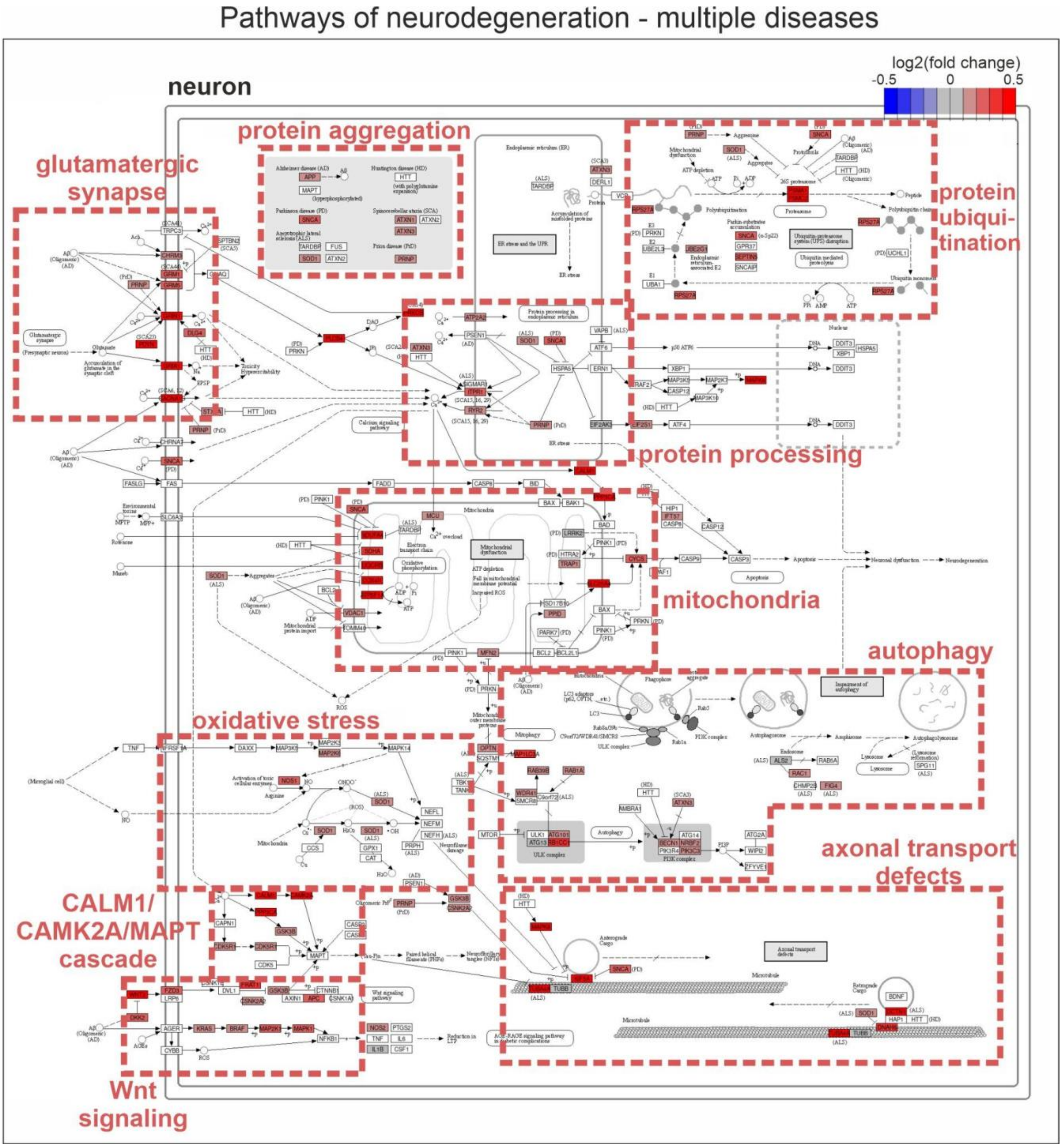


**Supplementary Fig. 5 Illustration of the KEGG pathway "Pathways of neurodegeneration - multiple diseases (hsa05022ID)".**

Overview of the KEGG pathway "Pathways of neurodegeneration - multiple diseases (hsa05022)" with genes colored by their log2 fold change in human female samples (as shown in Fig. 1h). This pathway contains parts related to the accumulation of protein aggregates, protein processing in ER/ER stress, protein ubiquitination (ubiquitin-proteasome system dysfunction), impaired glutamatergic synapse, impaired Wnt- and Calcium/CAMK2A/MAPT cascade (formation of neurofibrillary tangles), mitochondrial dysfunction, oxidative stress, autophagy impairment, and axonal transport defects. Adapted from the KEGG database (https://www.genome.jp/pathway/hsa05022).

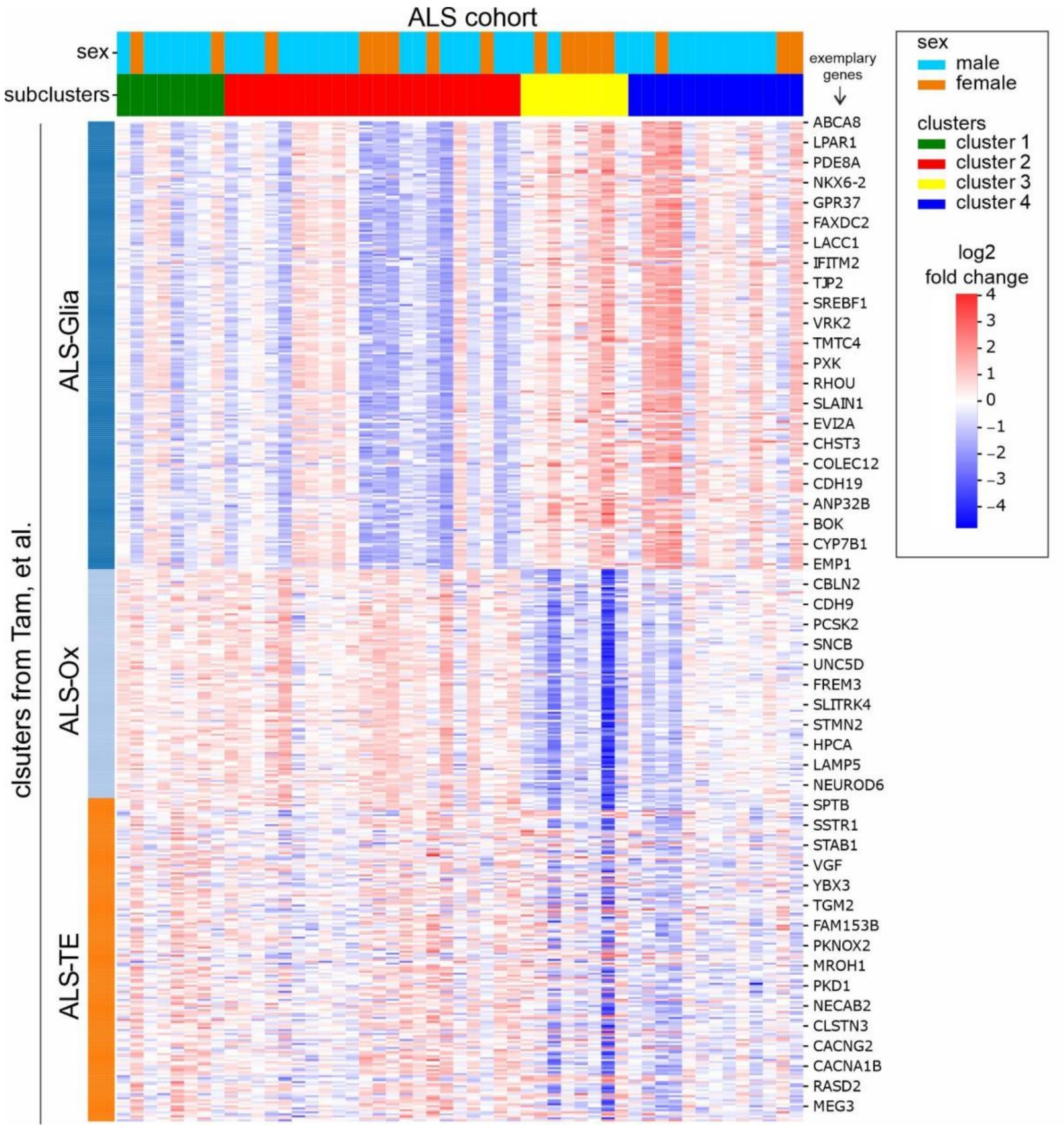


**Supplementary Fig. 6 Expression of genes involved in ALS subclusters by Tam et al. 2019.**

Genes were selected based on Supplemental file 4 (table S2A) from Tam et al. (2019)[1] grouped into ALS-TE (orange), ALS-Ox (light-blue) and ALS-Glia (dark-blue). Gene expression is vst-transformed and mean-variance scaled for display. The sample-subclusters are identified using pathway information as described in Fig 1. Only a representative number of gene names is shown.

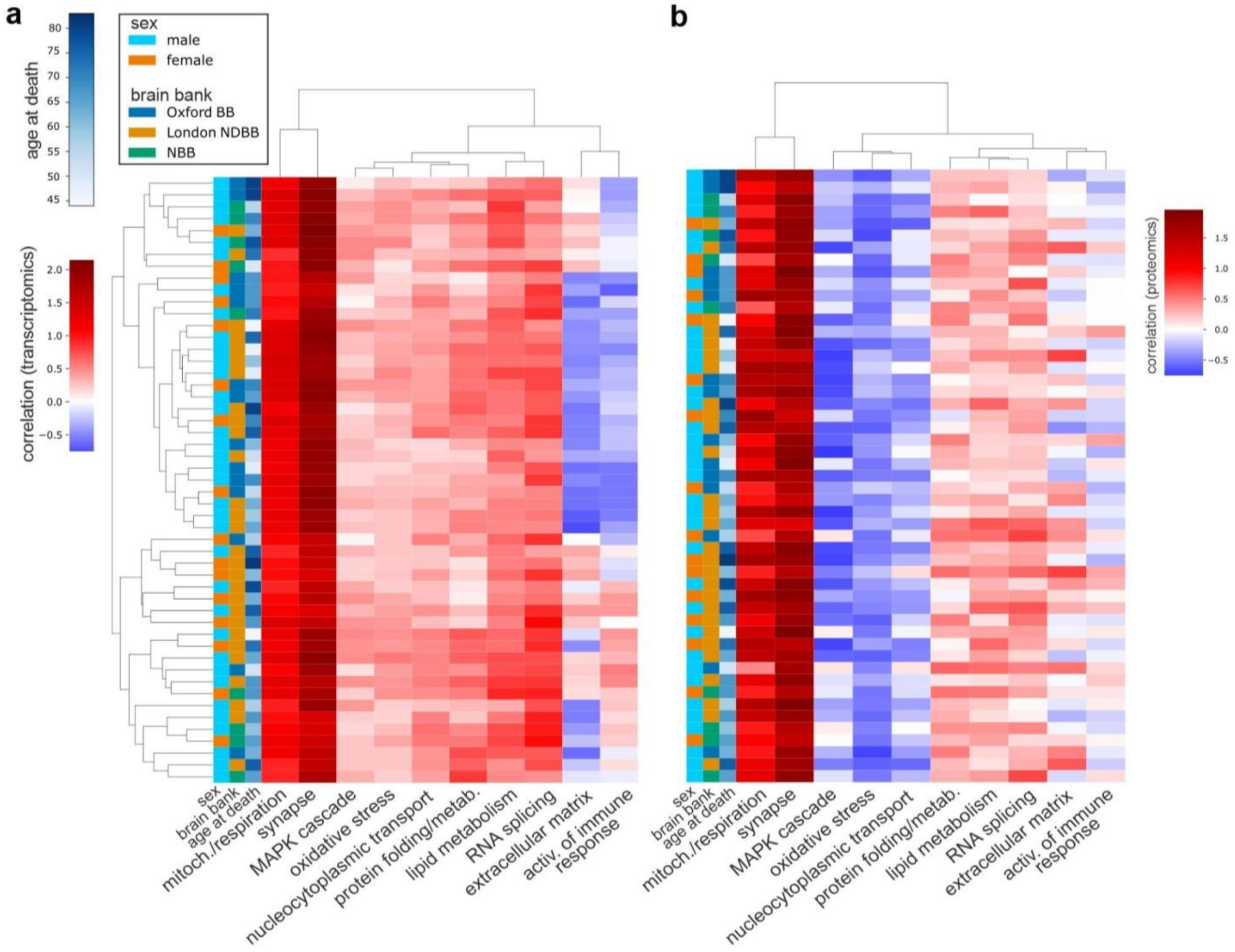


**Supplementary Fig. 7 Projection of transcriptomic subclusters onto proteomic data.** Heatmap depicts activity scores (calculated by Decoupler) of each selected pathway throughout the entire ALS human cohort in the transcriptomics analysis (**a**) and the proteomics analysis (**b**). Pathways are indicated on the y-axis and the ALS samples on the x-axis (top) along with the metadata, sex, brain bank and age at death.

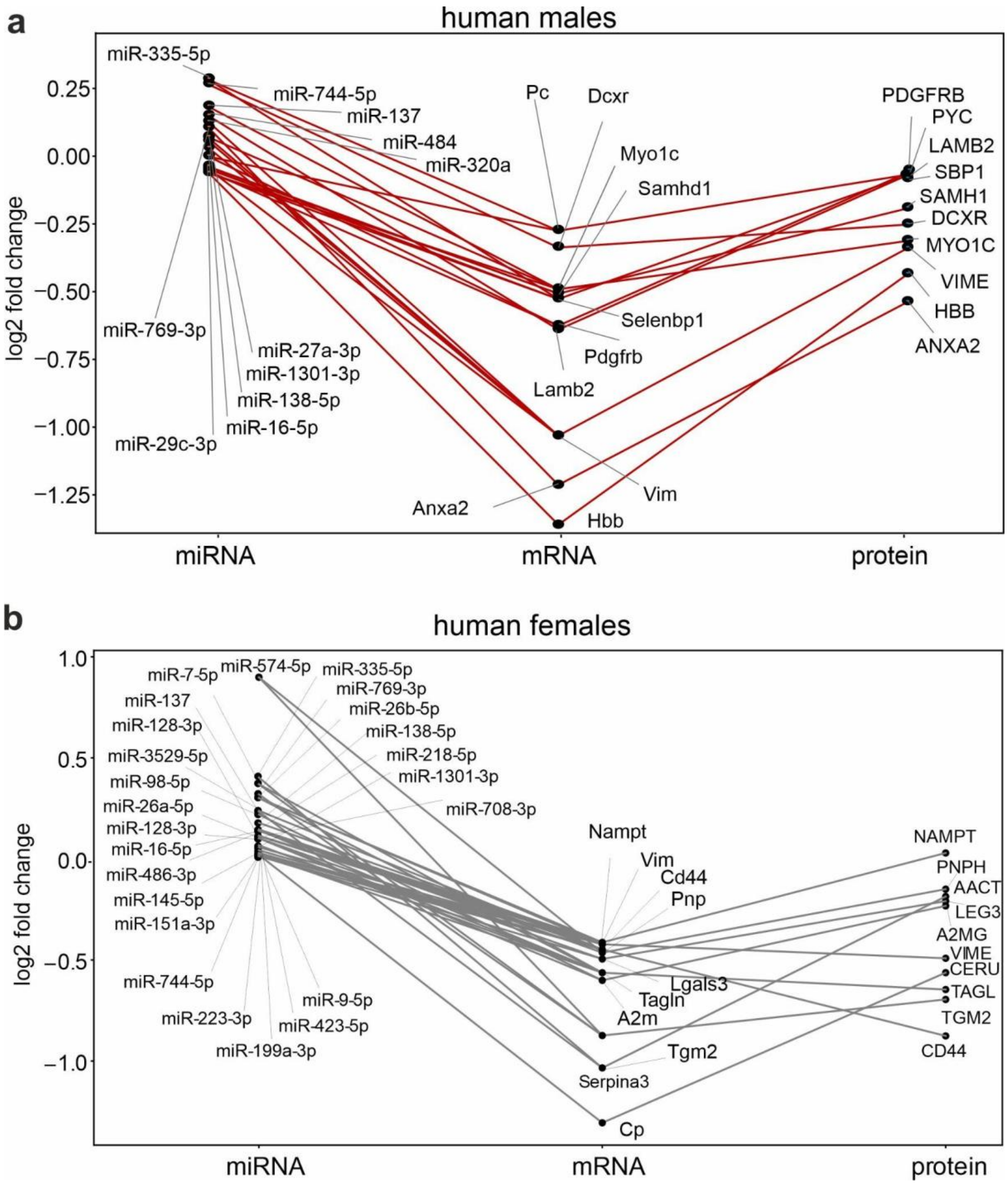


**Supplementary Fig. 8 Triplet-plots showing miRNA - mRNA - protein networks in human males and females.**

Schematic view of the miRNA - mRNA - protein - regulatory network for human males **(a)** and females **(b)**, for ALS vs. CTR comparisons. We selected significantly altered miRNAs based on the adjusted p-values < 0.05 for males. For females, no significant changes were observed (therefore displayed in grey), hence we depict the top 10 miRNAs based on fold change. The y-axes show the log2 fold change, with positive values indicating upregulation and negative values indicating downregulation. The x-axes show miRNA, mRNA and protein names. The connecting lines show miRNA-mRNA-protein networks. Only networks with opposite regulation between miRNA and mRNA (down-regulated transcripts) are shown. Networks consisting of mRNAs with a log2 fold change within the top 10 log2 fold changes of all mRNA are shown in red.

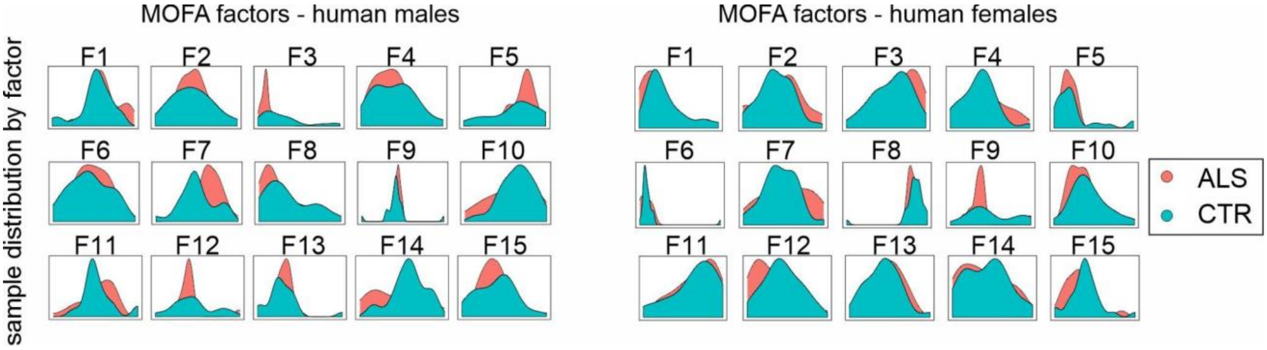


**Supplementary Fig. 9 Density plot for factors identified with MOFA.**

Density plots displaying the sample distribution by factor for male and female samples in the MOFA analysis. X-axis, MOFA factor values; y-axis, number of samples. Red: ALS, turquoise: CTR.

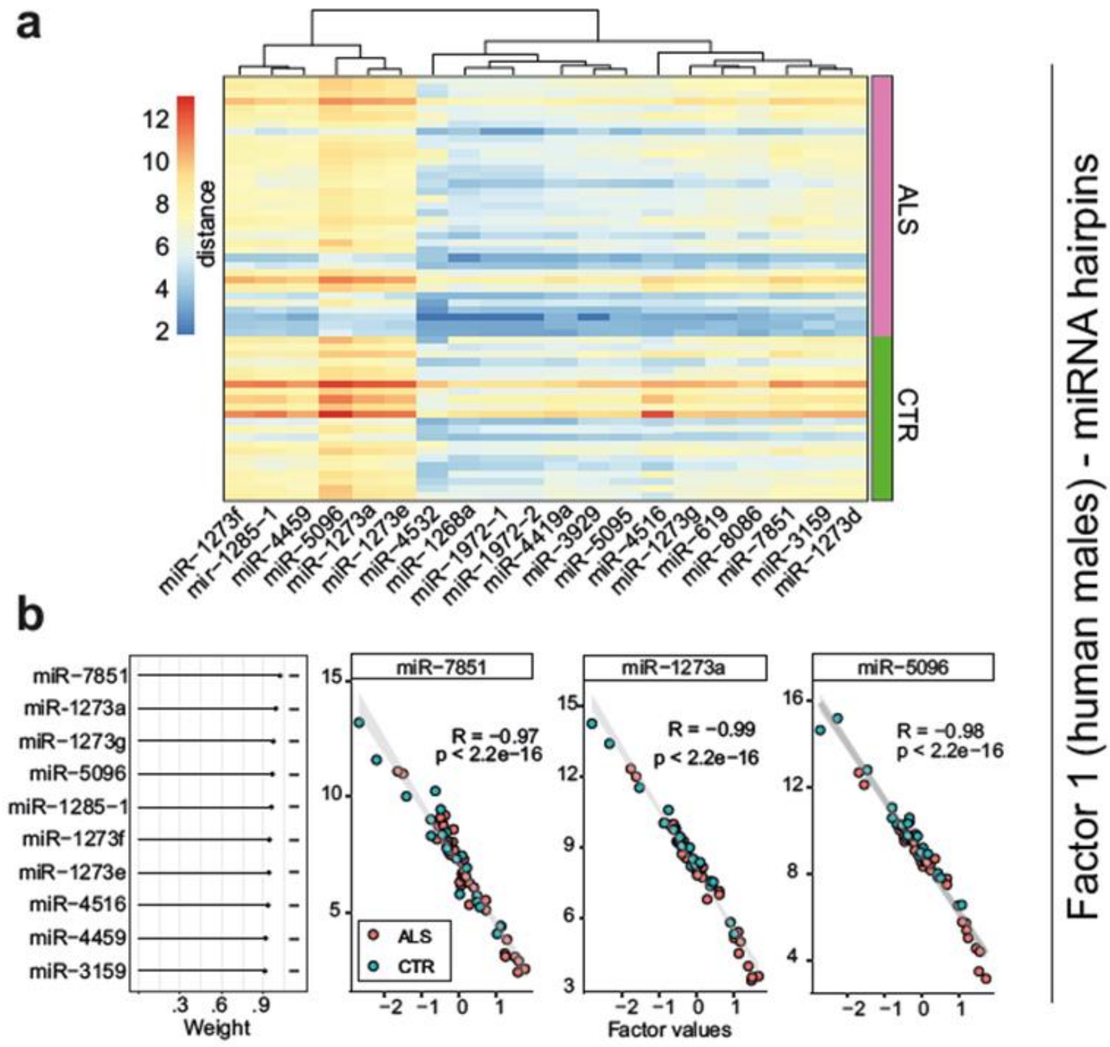


**Supplementary Fig. 10 Overview of MOFA factor 1 for male ALS patients.**

MOFA analysis of multiomic data from male human samples identifies molecular factors that contribute to the variability between samples. Overview of the representative factor 1, mainly driven by miRNA hairpins. **a.** Expression patterns of miRNAs that contribute most to factor 1. Color-coded heatmap based on the expression levels of miRNAs: blue, low expression; red, high expression. Heatmap dendrogram shows the hierarchical clustering of miRNAs based on their expression patterns. ALS samples, pink bar; CTR samples, green bar. **b.** MOFA weight plot for hairpin miRNAs in human males displaying the contribution of each miRNA to the MOFA model. Each line represents a miRNA, the x-axis shows the MOFA weight for that miRNA, reflecting its importance in the MOFA model. Scatter plots show the relationship between the expression of selected miRNAs for male samples and the factor values identified in factor 1. Red: ALS, turquoise: CTR.

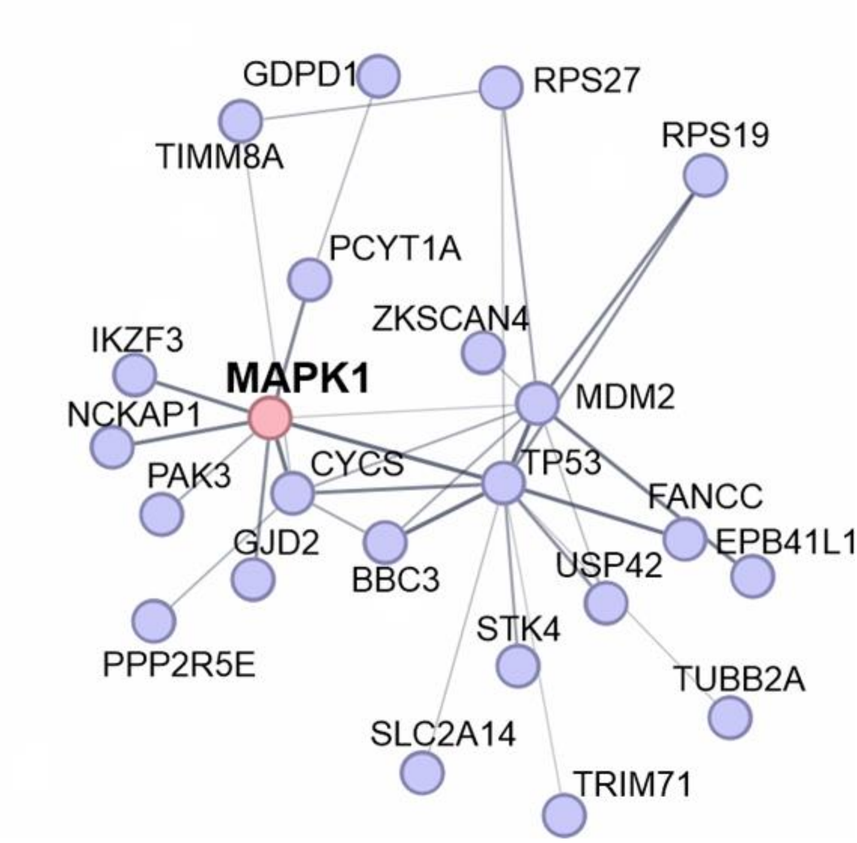


**Supplementary Fig. 11 Protein-protein-interaction network for the interactors of MAPK1 among the target genes of miR-1273f.**

Interactors of up to two nodes of distance are depicted.

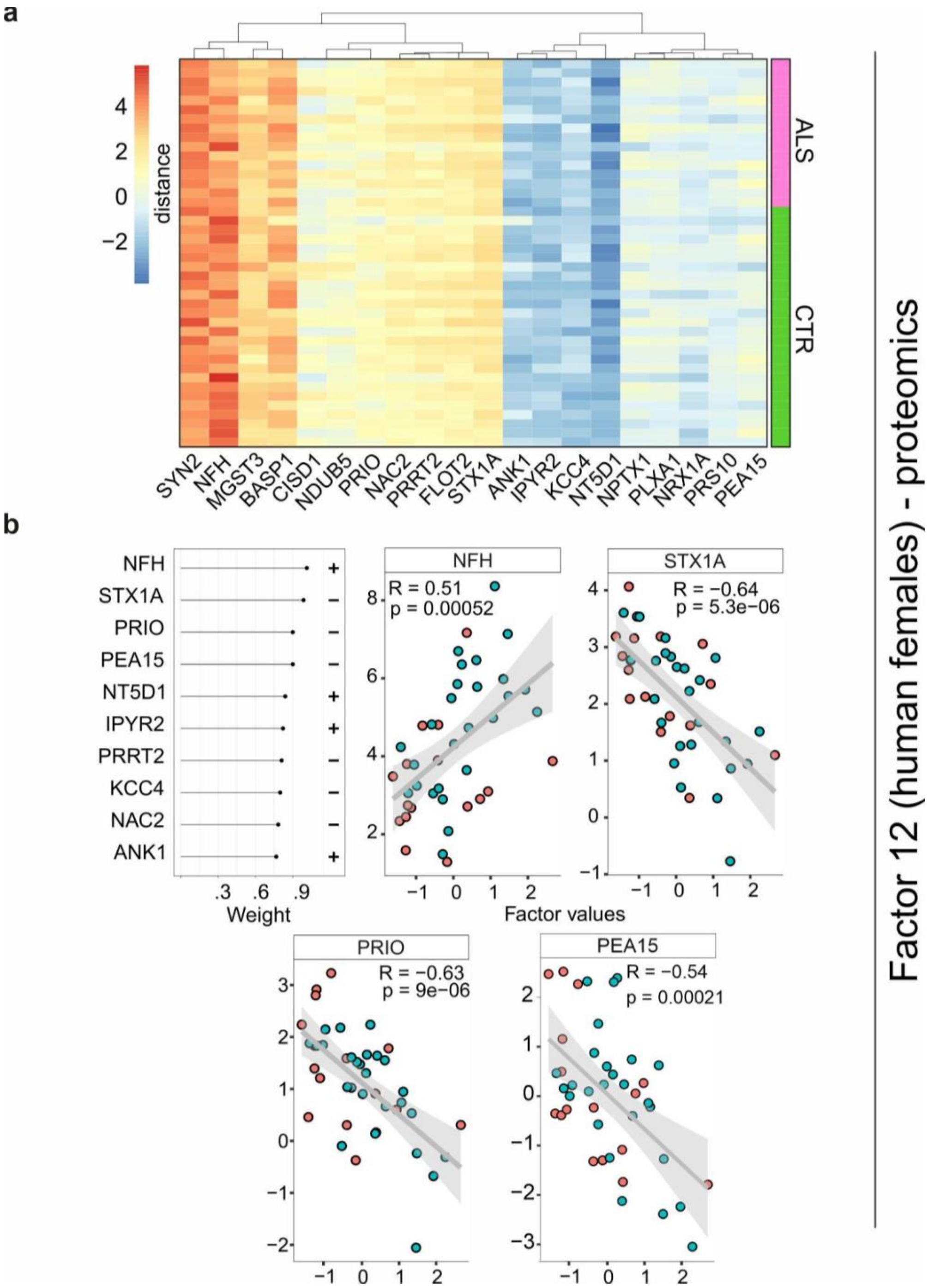


**Supplementary Fig. 12 Overview of MOFA factor 12 for female ALS patients.** Overview of the representative factor 12, which contributed most to the dichotomization of ALS and CTR samples (for females), mainly driven by proteomics. **a.** Expression patterns of proteins that contribute most to factor 12. Color-coded heatmap based on the abundance of proteins: blue, low abundance; red, high abundance. Heatmap dendrogram shows the hierarchical clustering of proteins based on their abundance patterns. ALS samples, pink bar; CTR samples, green bar. **b.** MOFA weight plot for proteins in human females displaying the contribution of each protein to the MOFA model. Each line represents a protein, the x-axis shows the MOFA weight for that protein, reflecting its importance in the MOFA model. Scatter plots show the relationship between the abundance of selected proteins for female samples and the factor values identified in factor 12. Red: ALS, turquoise: CTR.

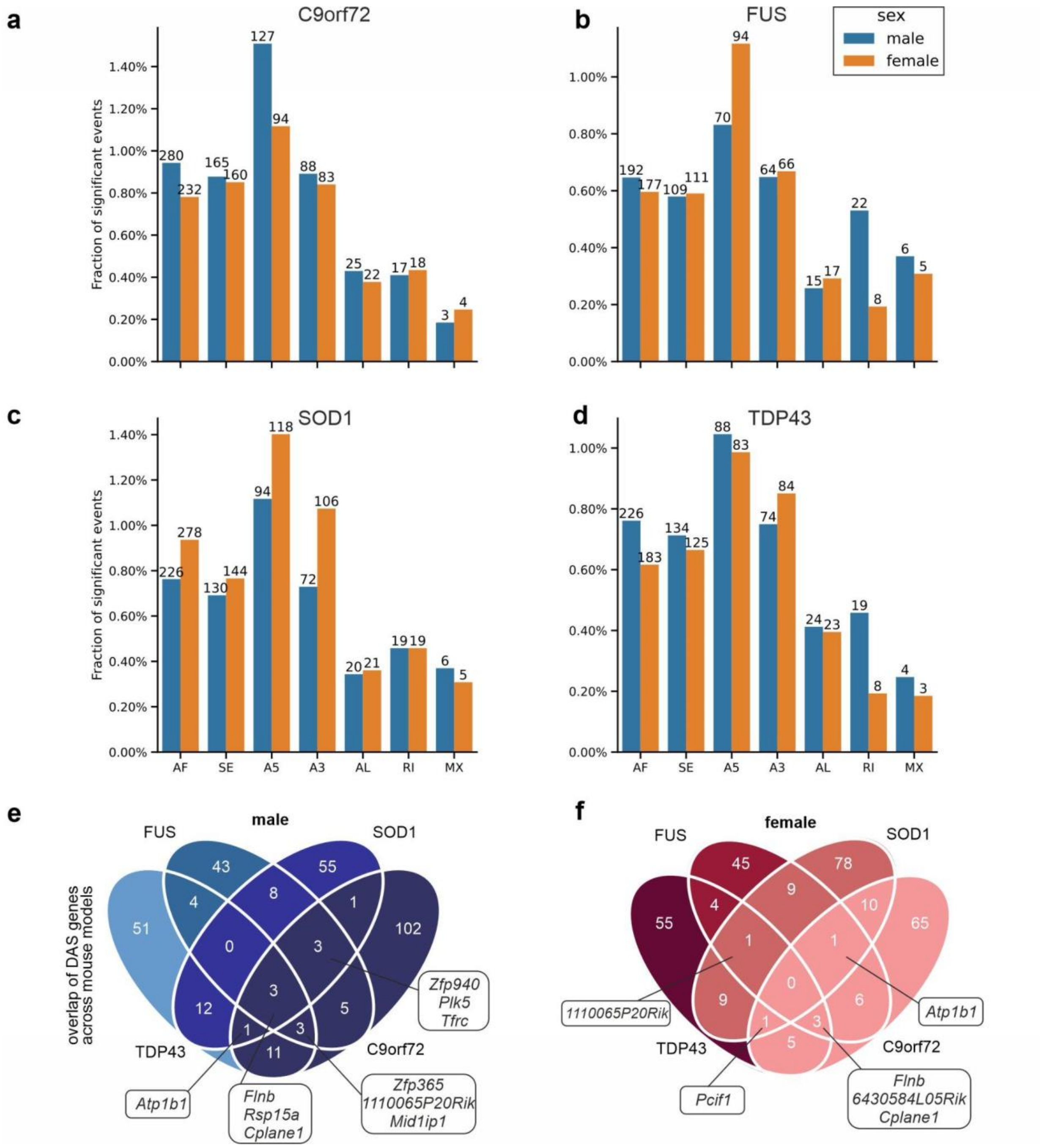


**Supplementary Fig. 13 Differential alternative splicing (DAS) analysis of the mouse models.**

**a-d.** Differential Alternative Splicing (DAS) analysis of male and female samples in four transgenic mouse models: C9orf72 **(a)**, FUS **(b)**, C9orf72 **(c)** SOD1, and TDP43 **(d)**. The following splicing events are considered: alternative exon (AE), skipped exon (SE), alternative 5' splice site (A5), alternative 3' splice site (A3), alternative last exon (AL), retained intron (RI) and mutually exclusive exon (MX) events. Each event is represented by a separate bar, with the height of the bar representing the fraction of significant events in ALS vs. CTRL. Events with significant differential splicing in males (blue) and females (orange). **e-f.** Venn diagrams showing the overlap of DAS genes in the studied mouse models compared in ALS vs CTR. Each circle represents the number of genes with significant DAS in each model. Overlapping regions between the circles represent the number of genes with significant DAS in more than one model.

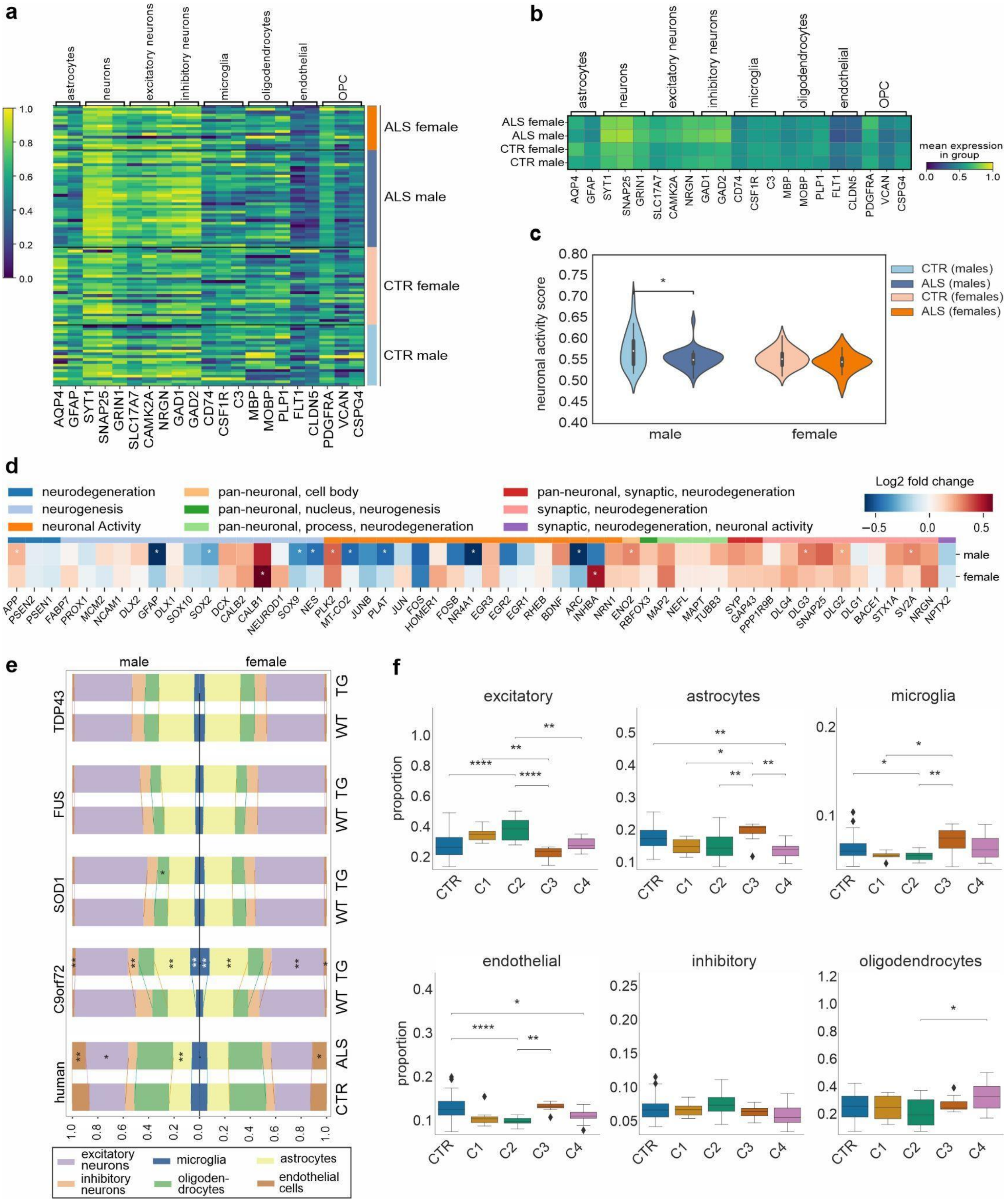


**Supplementary Fig. 14 Cell-type deconvolution.**

**a**. Heatmap representing the expression of cell-type-specific marker genes in human samples, each column representing one cell type (astrocytes, neurons, microglia, excitatory neurons, inhibitory neurons, oligodendrocytes, endothelial cells, and oligodendrocyte-precursor cells [OPC]), each row representing one sample. Marker genes shown on x-axis. ALS males, light blue; ALS females, dark blue; control males, light orange; control females, dark orange. **b.**

Heatmap showing mean expression of selected neuronal cell type marker genes in ALS, CTR, male and female. **c.** Violin plot displaying the distribution of neuronal activity scores in ALS and CTR samples by sex estimated using NEUROeSTIMator (https://research-git.uiowa.edu/michaelson-lab-public/neuroestimator). ALS, brown; CTR, blue. * p <= 0.05 by 2-tailed Student's T-test. **d.** log2 fold change of expression of genes representative of different neuronal compartments or processes. * p <= 0.05 by Wald test. e. Mean cell type fractions estimated on mouse models and human mRNA samples. The lines connecting bars indicate changes in the mean cell type fractions from CTR to ALS. ** p < 0.01, * p < 0.05 by independent T-test. f. Boxplots showing estimated cell-type proportions across the four clusters C1-C4 identified in the ALS samples (Fig. 1f). * p <= 0.05, **p <= 0.01, **** p <= 0.0001 by ANOVA with post-hoc Tukey's range test for pairwise group-mean comparisons.

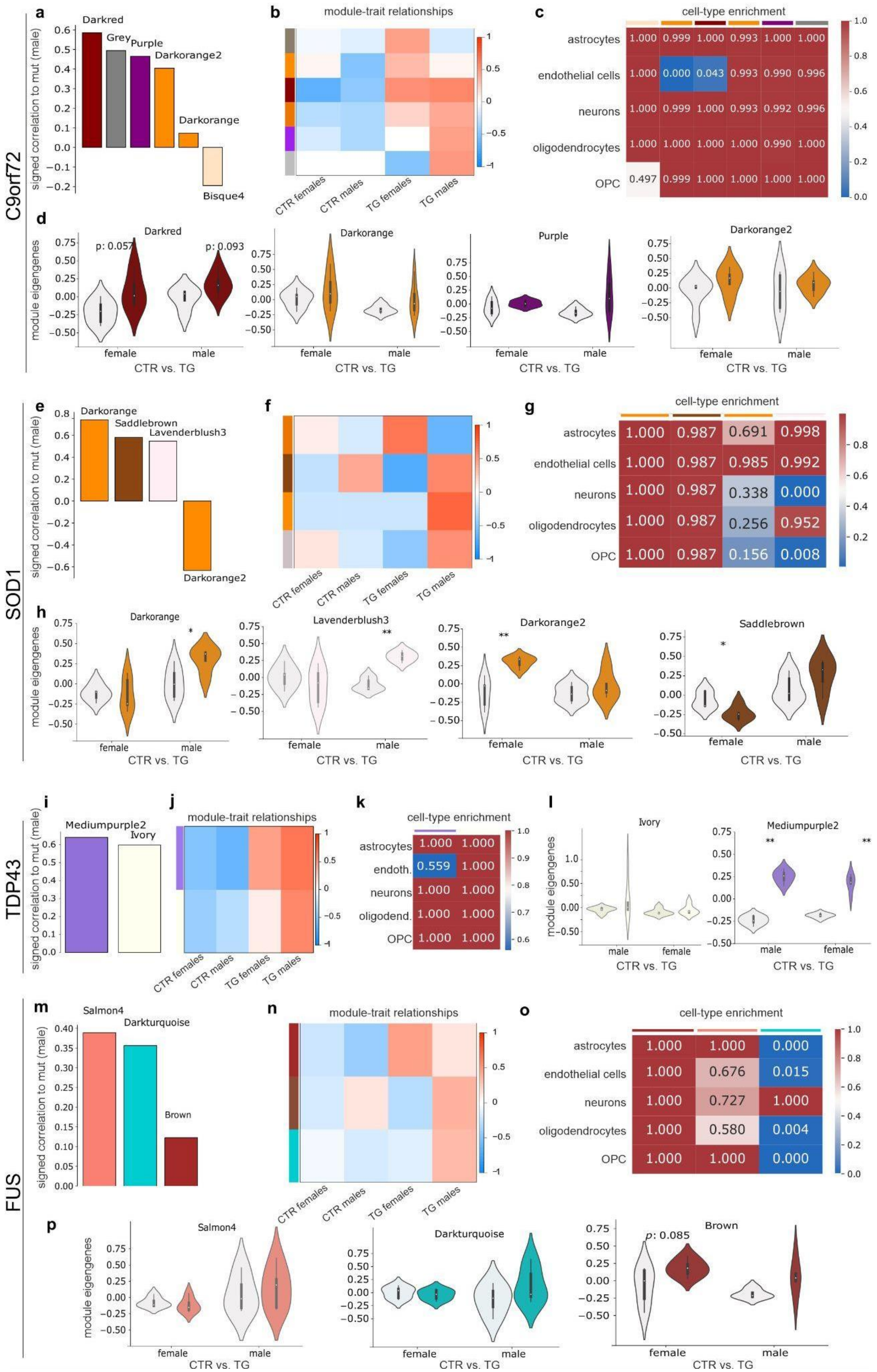


**Supplementary Fig. 15 Summary of WGCNA for mouse models.**

Figures summarizing the mRNA WGCNA modules inferred for males and females between TG and CTR. **a-d WGCNA results for the C9orf72 model. a.** The Darkred, Darkorange, Darkorange2 and Purple modules were significantly and positively associated with TG males and females. **b.** Pearson's correlation between module eigengenes and sample groups for the C9orf72 model. Sex: M, F; condition: TG, WT. ** $p < 0.01$, * $p < 0.05$ by Student's t-test, for all models and comparisons. **c.** Heatmap showing adjusted p-values associated with the cell type enrichment on gene sets of each of the four modules. The background gene set was taken from Pangalo DB for all models. **d.** Violin plots showing the distribution of module eigen-genes in male and female mice and between TG and WT. **e-h WGCNA results for the SOD1 model**. **e.** The Darkred, Darkorange, Darkorange2 and Purple modules were significantly and positively associated with TG males and females. **f.** Pearson's correlation between module eigengenes and sample groups for the SOD1 model. **g.** Heatmap showing adjusted p-values associated with the cell type enrichment on gene sets of each of the four modules. **h.** Violin plots showing the distribution of module eigengenes in male and female mice and between TG and WT. **i-l WGCNA results for the TDP43 model. i.** The Ivory module was significantly and positively associated with TG males and females. **j.** Pearson's correlation between module eigengenes and sample groups for the TDP43 model. **k.** Heatmap showing adjusted p-values associated with the cell type enrichment on gene sets of each of the four modules. **l.** Violin plots showing the distribution of module eigengenes in male and female mice and between TG and WT. **m-p WGCNA results for the FUS model. m.** The Salmon4, Darkturqoise and Brown modules were significantly and positively associated with males and females. **n.** Pearson's correlation between module eigengenes and sample groups for the FUS model. **o.** Cell type enrichment on gene sets of each of the four modules. **p.** Violin plots showing the distribution of module eigengenes in male and female mice and between TG and WT. TG, transgenic mice; WT, wild-type mice.

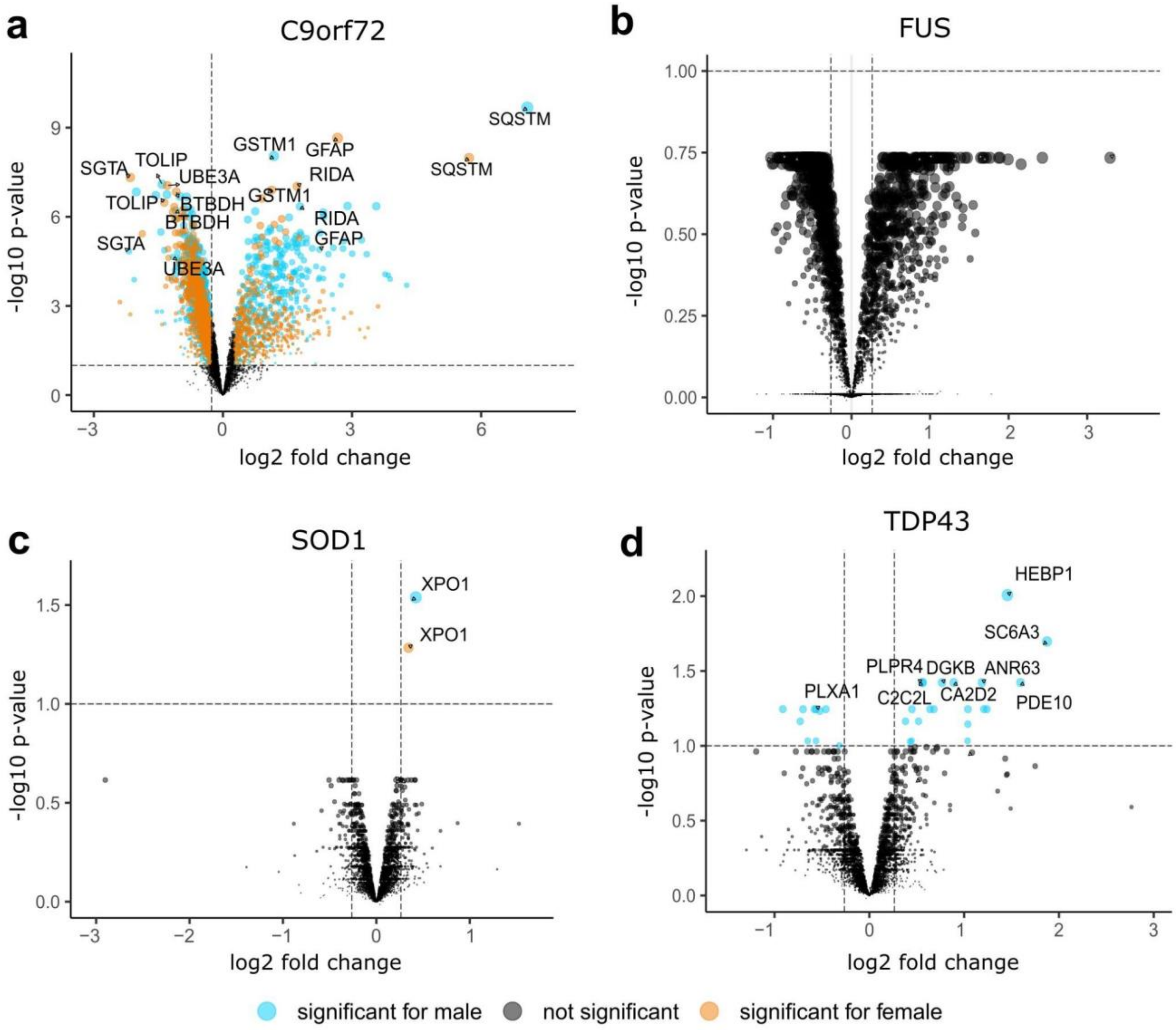


**Supplementary Fig. 16: Differential expression of proteins in mouse models.**
Volcano plots displaying the results of the proteomics analysis in ALS mouse models, comparing C9orf72 **(a)**, FUS **(b)**, SOD1 **(c)**, and TDP43 **(d)** models versus controls, stratified by sex. Each dot on the plot represents one protein. The x-axis displays the log2 fold change in protein expression between transgenic mice and wild-type litter-mates. Positive values indicate upregulation in ALS. Y-axis displaying the negative log10 p-value, with higher values indicating more significant differences in expression. The horizontal dashed line represents the significance threshold, p-value = 0.1, and the vertical dashed lines represent the threshold for log2 fold change (±1.5). Males, blue circles; females, orange circles.

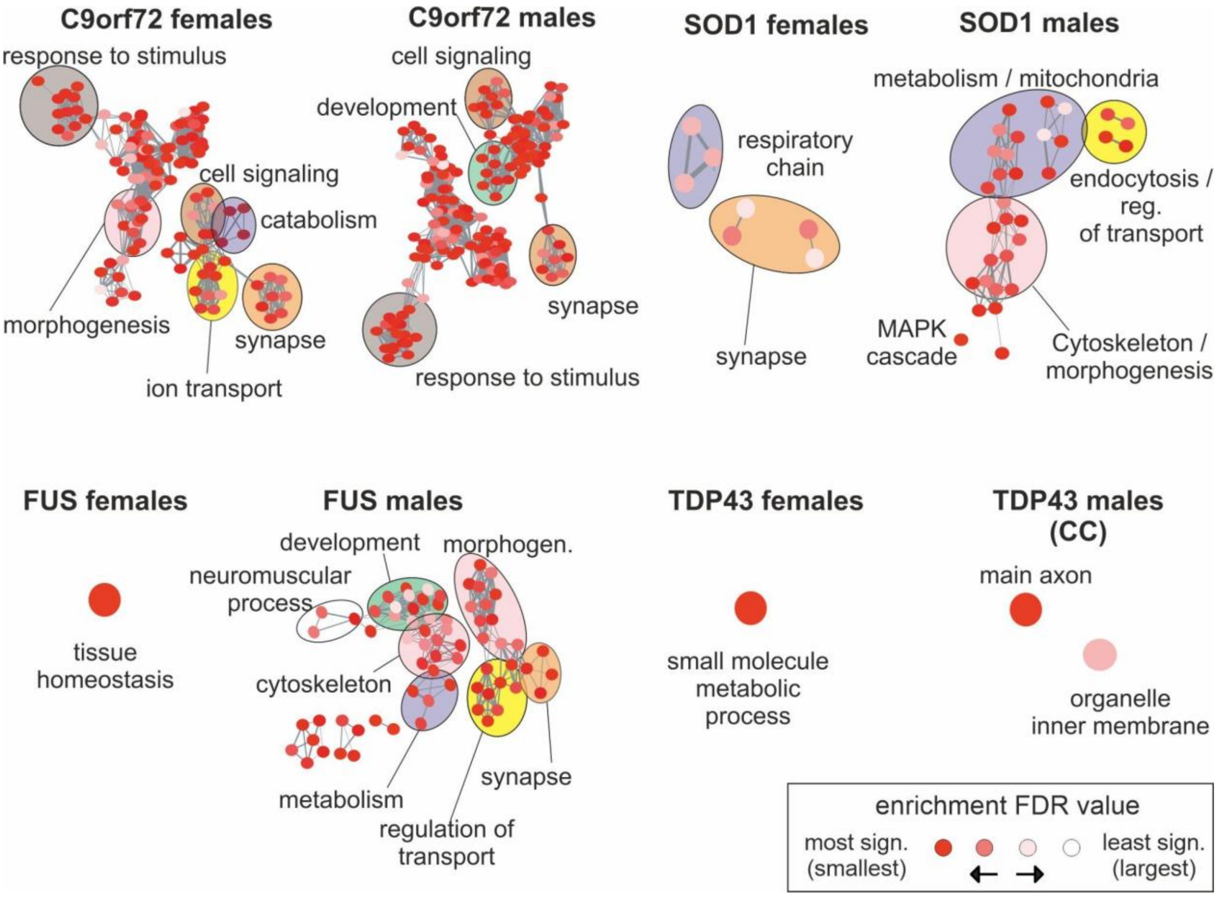


**Supplementary Fig. 17: REVIGO clustering and sample correlation for mouse proteomics.**

REVIGO-based summary of proteomics gene set enrichment results for the mouse samples. The plots summarize the functional similarity of the proteins by reducing redundant GO-BP terms and clustering the remaining non-redundant terms. Each area of the plot represents a cluster of similar GO BP terms. The size of the circles represents the number of genes in the GO-BP terms, and the color of each node represents the enrichment FDR values.

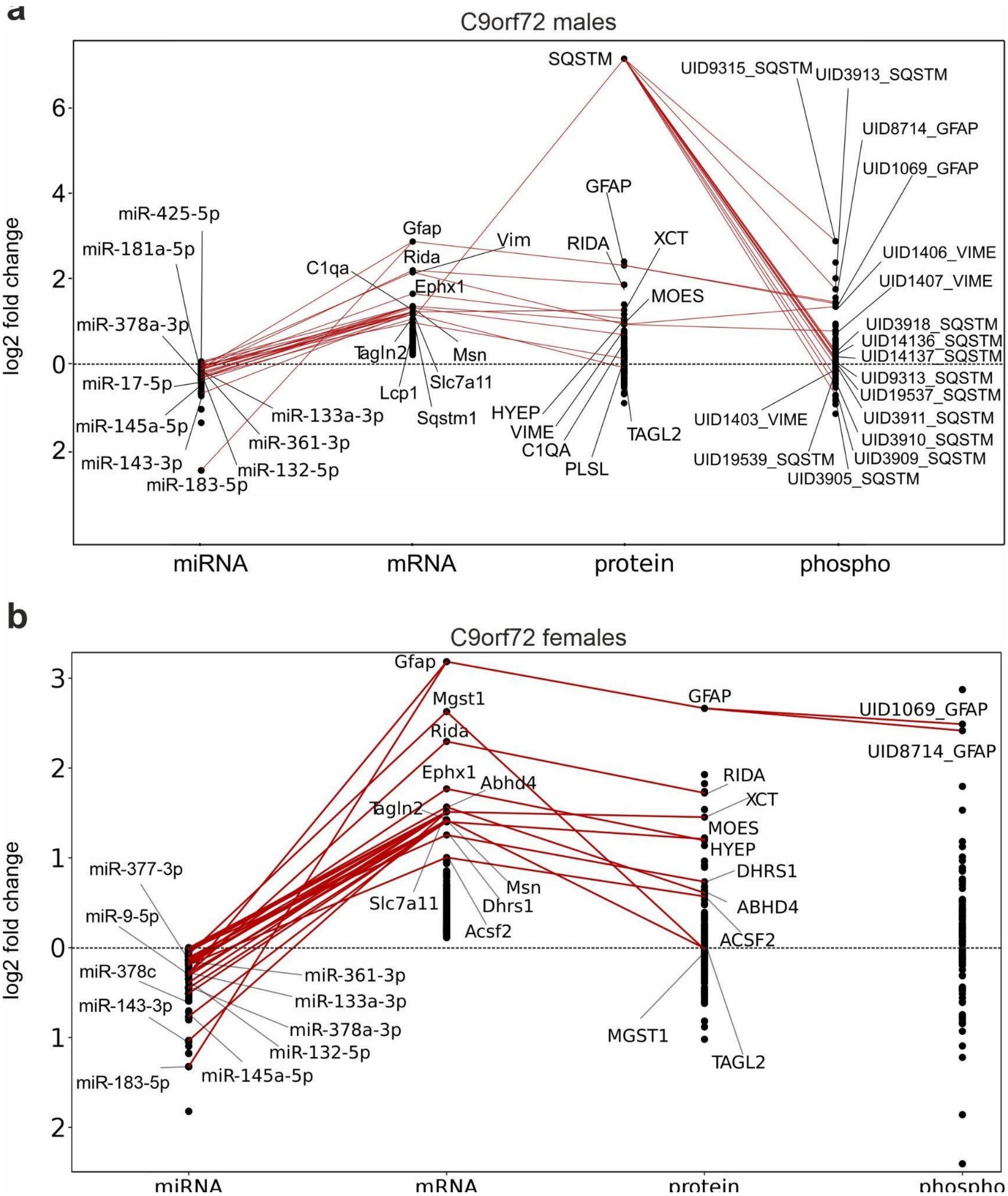


**Supplementary Fig. 18: Quadruple plots showing miRNA - mRNA - protein - phosphoprotein networks in the C9orf72 model.**

Schematic view of the miRNA - mRNA - protein - phosphoprotein regulatory network for the C9orf72 mouse model for males (**a**) and females (**b**). A cutoff of 0.05 was used on the adjusted p-values on mRNA log2 fold change. The top 10 most downregulated miRNAs with valid interactors were considered. The x-axis represents the different stages of regulation: miRNA, mRNA, protein, and phosphoprotein. The y-axis represents the abundance (log2 fold change) of each entity. Full integrative results for all cohorts are provided in Supplementary Table 10.

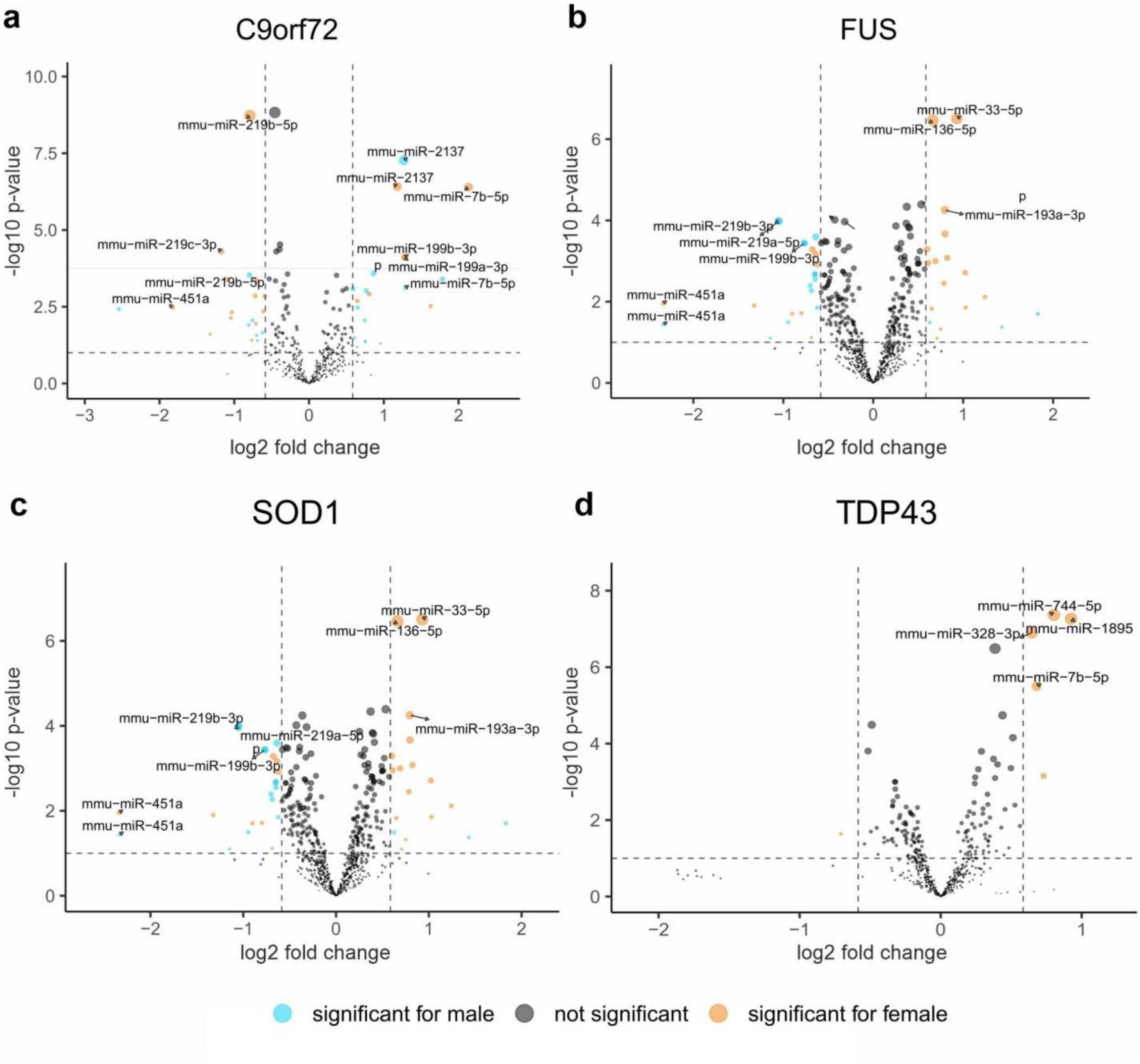


**Supplementary Fig. 19. Differential expression of miRNA in mouse models.**

**a-d** Volcano plots displaying the results of miRNAomics analysis in ALS mouse models, comparing transgenic animals in C9orf72 **(a)**, FUS **(b)**, SOD1 **(c)**, and TDP43 **(d)** models versus wild-type litter-mates, stratified by sex. Each dot on the plot represents one miRNA. The x-axis displays the log2 fold change in miRNA expression between transgenic animals and wild-types. Positive values indicate upregulation in transgenics. The y-axis displays the negative log10 p-value, with higher values indicating more significant differences in expression. The horizontal dashed line represents the significance threshold, p-value = 0.1, and the vertical dashed lines represent the threshold for log2 fold change (±1.5). Males, blue; females, orange.

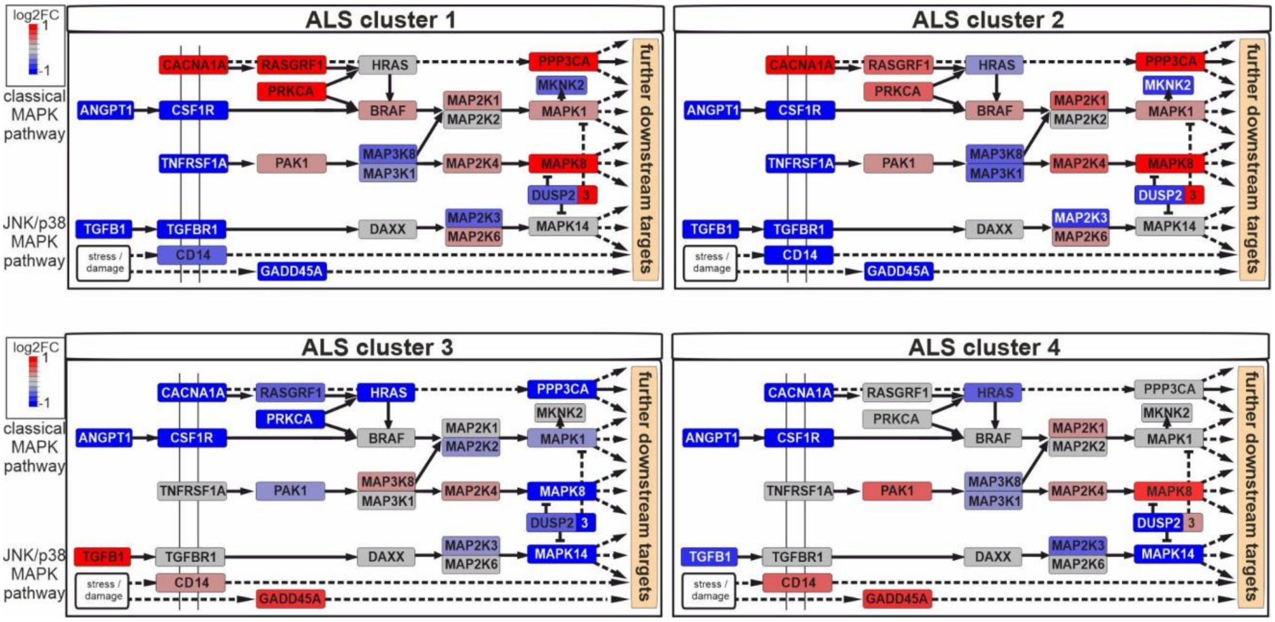


**Supplementary Fig. 20 Differential regulation of MAPK pathways in human ALS patient subclusters.**

Analysis of human ALS clusters revealed consistent changes in MAPK signaling across different subclusters. The panels show the classical MAPK pathway that is found activated in C1 and C2, and downregulated in C3. The JNK and p38 MAPK pathways are downregulated in C1 and C2, and activated in C3. C4 shows a more subtle deregulation in those pathways, with similarities to C3 especially for upstream molecular players. The panels depict log2 fold change values for transcriptomics in colors.

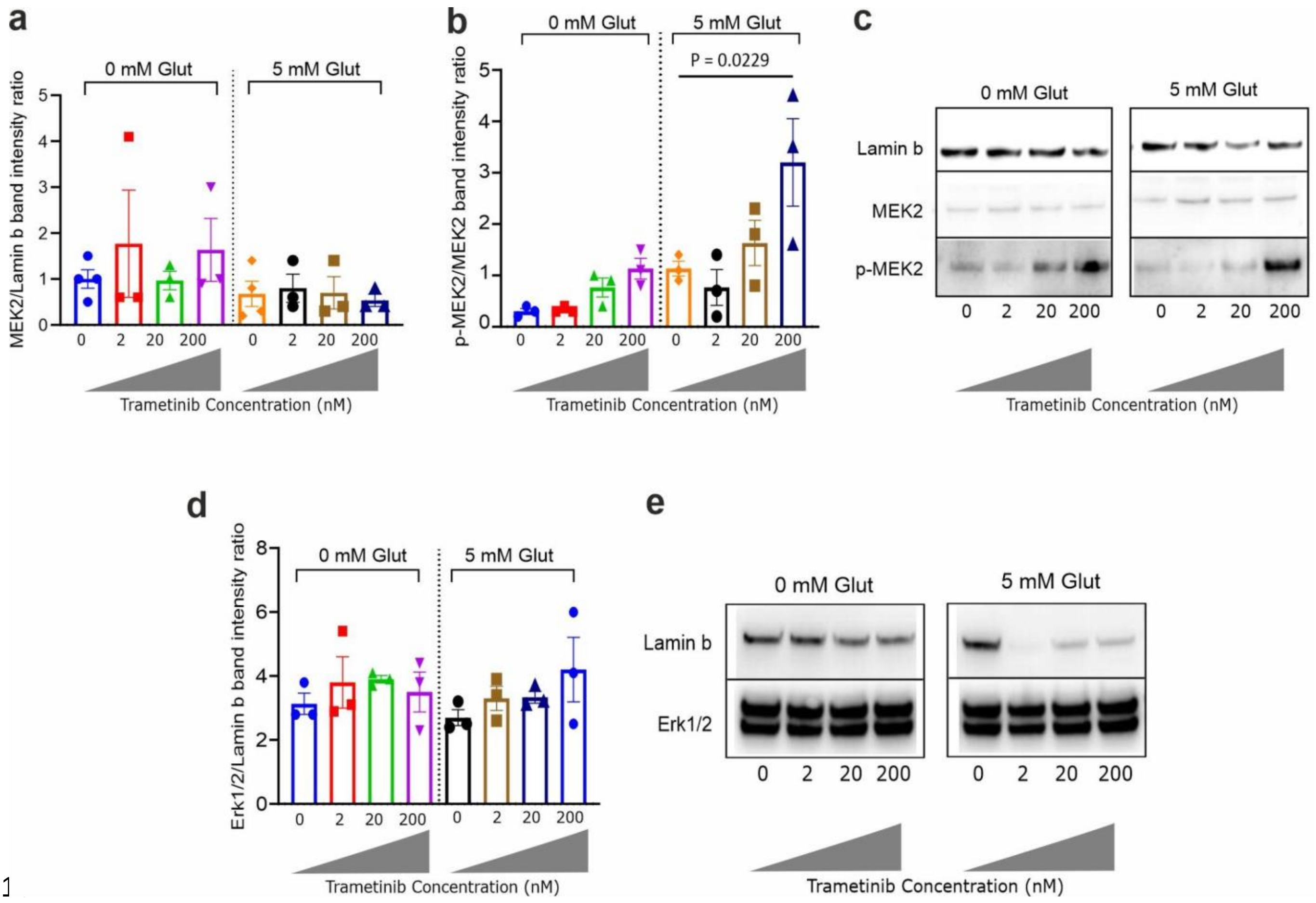


**Supplementary Fig. 21 Western blot analysis of trametinib effects on MEK2, ERK1/2 and phospho-MEK2 with and without glutamate treatment in primary neuronal cultures.**

Western blot quantification of MEK2 normalized on lamin b **(a)** or phospho-MEK2 normalized on MEK2 **(b)** for both 0 mM Glutamate (left) and 5 mM Glutamate (right) treatment conditions for different trametinib concentrations (0 nM, 2 nM, 20 nM, 200 nM). **c.** Representative western blots. **d.** Western blot quantification of ERK1/2 normalized on lamin b for both 0 mM Glutamate (left) and 5 mM Glutamate (right) treatment conditions for different trametinib concentrations (0 nM, 2 nM, 20 nM, 200 nM). **e.** Representative western blots. Data are represented as the mean ± SEM of at least 3 different cultures and tested by ordinary one-way ANOVA, p-value as indicated.

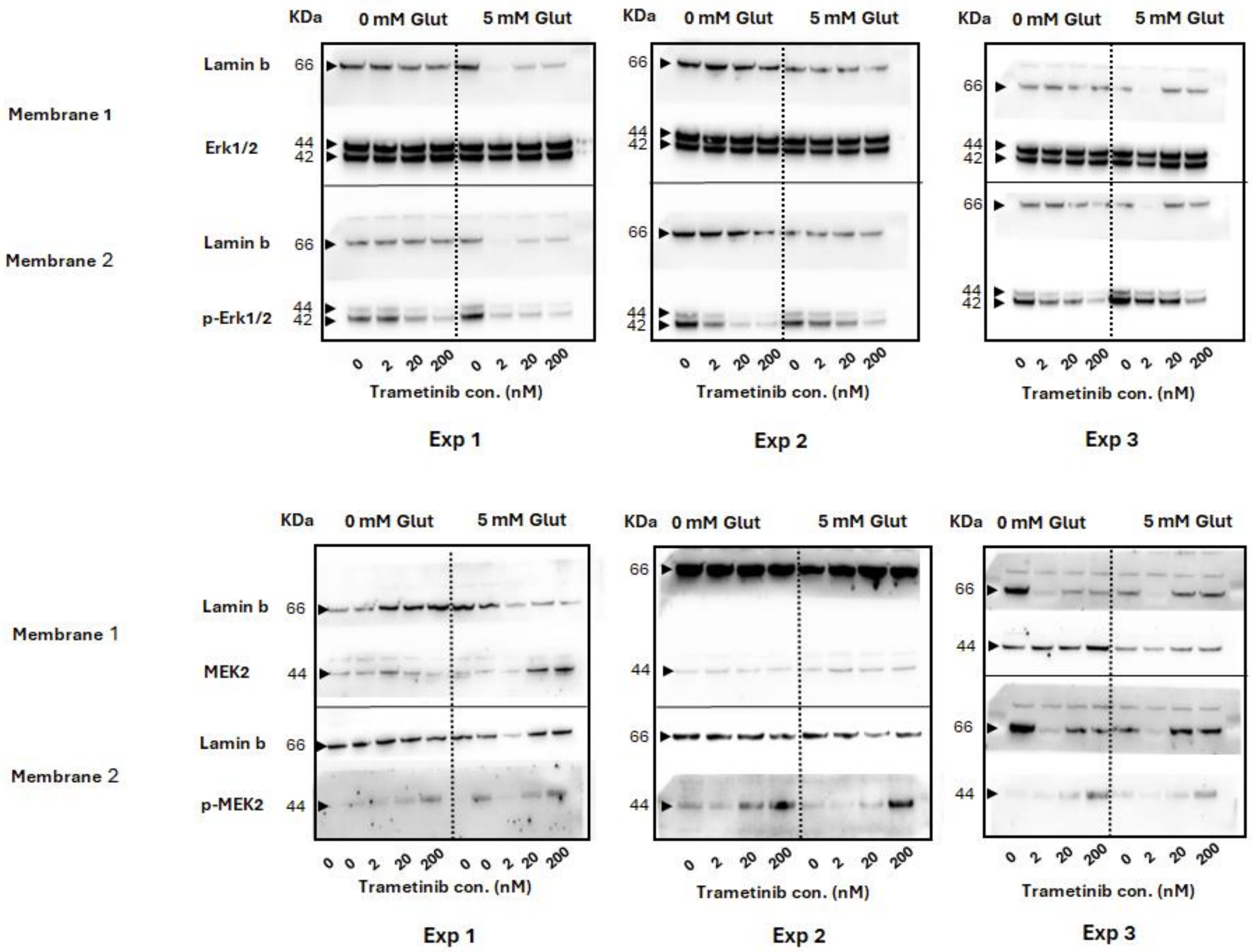


**Supplementary Fig. 22 Western blot experiments - full membranes.**
Three independent experiments for the assessment of Erk1/2 and MEK2 protein expression levels and their phosphorylation levels after treatment with different concentrations of trametinib, both with and without the presence of glutamate. To optimize resource and time utilization, membranes were precisely cut around the protein size and subsequently exposed to their respective antibodies.

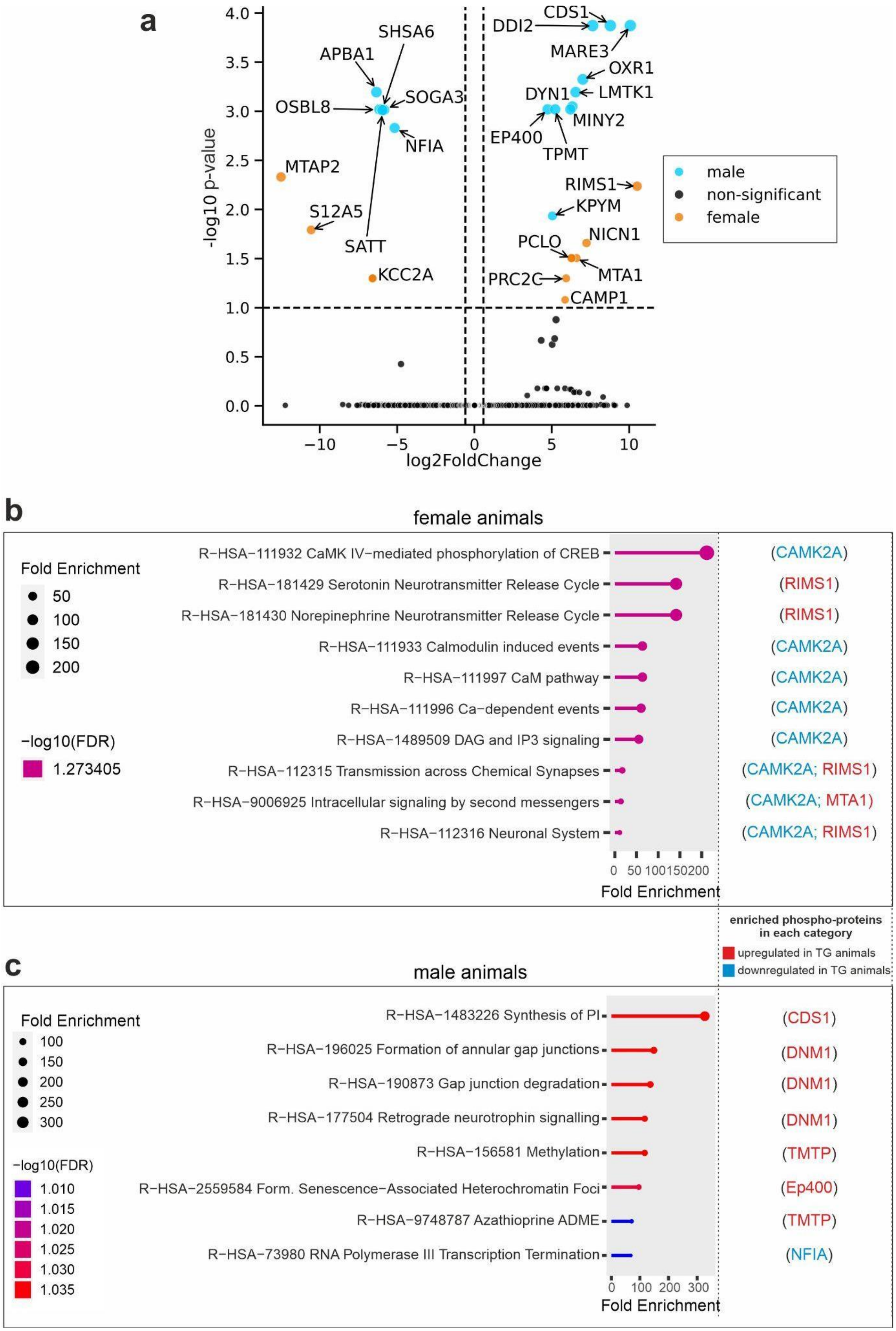


**Supplementary Fig. 23 Phospho-proteomics experiments for SOD1 mice treated with trametinib.**

**a** Volcano plot showing phospho-proteomics results from prefrontal cortex of SOD1 mice treated with trametinib (female: n=4; male: n=5) or vehicle (female: n=3; male: n=4). In total, 9 phosphoproteins were differentially regulated in females (6 up and 3 down) and 15 in males (10 up and 5 down) (padj = 0.1) following trametinib treatment. **b-c** Enrichment analyses for differentially regulated phospho-proteins for females **(b)** and males **(c).** The ShinyGO 0.80 curated database for Reactome (http://bioinformatics.sdstate.edu/go/) was used (enrichment FDR cutoff = 0.1).

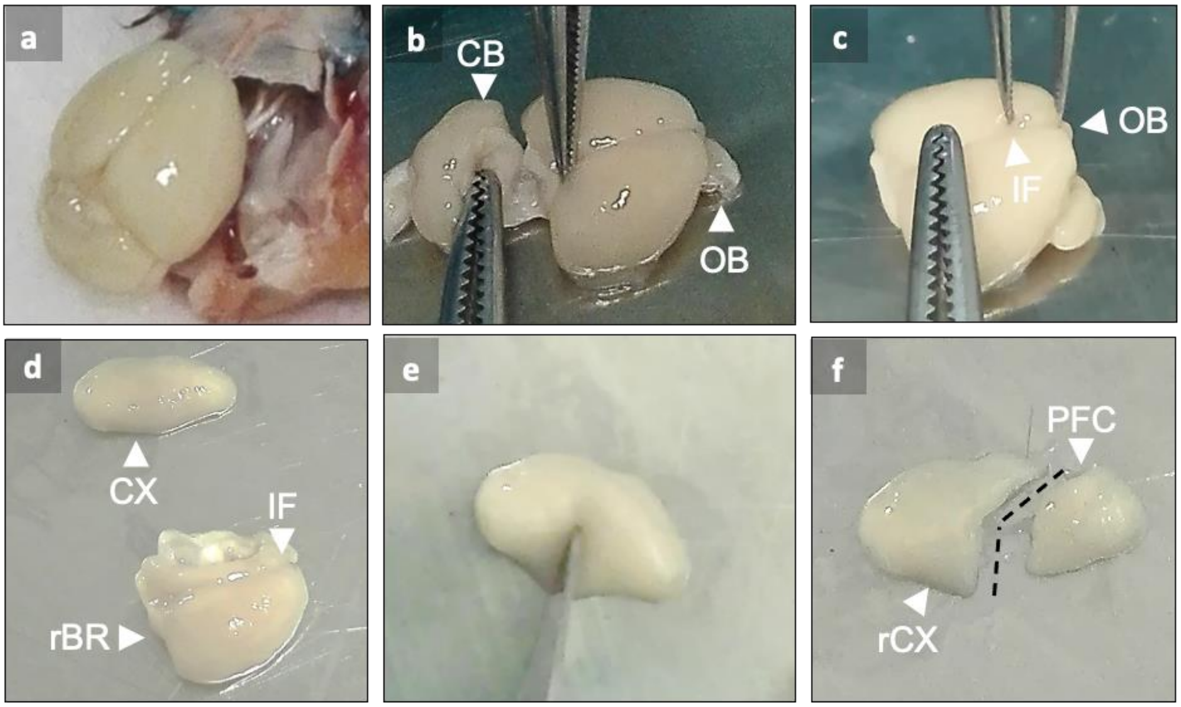


**Supplementary Fig. 24 Isolation of prefrontal cortex from ALS mouse models.**

Representative images of the procedure to isolate the prefrontal cortex (PFC) from mice. Once mice were perfused with ice-cold PBS to eliminate blood contamination, the brain was removed from the skull **(a)**. Next, the olfactory bulb (white arrow, OB) and cerebellum (white arrow, CB) were dissociated from the brain **(b)**. The cortex was lifted from the rest of the brain (white arrow, rBR) by using tweezers with fine tips, starting from the OB along the interhemispheric fissure (white arrow, IF) **(c)**. A perpendicular incision was made in the middle of the cortex **(e)** and with a 135° diagonal incision PFC was removed (white arrow) from the rest of the cortex (white arrow, rCX) **(e-f)**.
